# Intrahepatic reporter assay reveals leaky somatic blockade of L1 retrotransposition in mice

**DOI:** 10.64898/2026.02.12.705536

**Authors:** May Raya, Réka Karkas, Olivia Orsolya Verebi, Ede Migh, Gergely Imre, Kitti Vecsernyés-Nagy, Katinka Szalmási, Anna Georgina Kopasz, Dominik Sándor Kocsis, Péter Kálmán, Zoltán Lipinszki, Farkas Sükösd, Wenfeng An, Jef D Boeke, István Nagy, Péter Horváth, Andrea Mátés-Nagy, Lajos Mátés

**Affiliations:** Institute of Genetics, HUN-REN Biological Research Centre, Szeged, Hungary; Doctoral School of Multidisciplinary Medical Sciences, University of Szeged, Szeged, Hungary; Synthetic and Systems Biology Unit, Institute of Biochemistry, HUN-REN Biological Research Centre, Szeged, Hungary; Doctoral School of Biology, University of Szeged, Szeged, Hungary; ATGandCo Biotechnology Ltd, Mórahalom, Hungary; Department of Pathology, Péterfy Sándor Street Hospital, Budapest, Hungary; Department of Pharmaceutical Sciences, South Dakota State University, Brookings, USA; Institute for Systems Genetics and Department of Biochemistry and Molecular Pharmacology, NYU School of Medicine, New York, NY 10016 USA; Department of Biomedical Engineering, NYU Tandon School of Engineering, Brooklyn NY 11201; Seqomics Biotechnology Ltd, Mórahalom, Hungary; Single-Cell Technologies Ltd, Szeged, Hungary; Institute of AI for Health, Helmholtz Zentrum München, Neuherberg, Germany; Institute for Molecular Medicine Finland (FIMM), University of Helsinki, Helsinki, Finland

**Author notes:** **corresponding author:** Lajos Mátés. These authors contributed equally to this work (May Raya, Réka Karkas).

## Abstract

Long interspersed element-1 (LINE-1, L1) retrotransposition has long been proposed to occur in somatic tissues, yet direct experimental evidence distinguishing adult somatic events from early embryonic insertions has remained limited. Here we establish an intrahepatic L1 reporter assay that enables immunohistochemical detection and quantitative analysis of L1 retrotransposition *in vivo*. Using autonomous and non-autonomous L1 reporter variants, we demonstrate clearly detectable somatic L1 activity in the mouse liver. Comparative analysis of L1 activity in liver tissue and tumor-derived cell culture reveals that tumor cells preferentially restrict L1 at early regulatory stages, consistent with epigenetic control, whereas downstream defence mechanisms are comparatively permissive. In contrast, normal liver tissue shows stronger restriction at later stages of the L1 life cycle. Together, our results provide direct experimental evidence for somatic L1 retrotransposition *in vivo* in adult liver and reveal distinct regulatory strategies that shape L1 activity in tumor versus normal somatic cells.

**Teaser:** Genome destabilizing L1 retrotransposon activity is present in somatic tissues, where it likely contributes to cancer development.

## Introduction

The incidence of cancer has significantly increased worldwide in recent years (*1*), partly due to an ageing population (*2*), but also due to increased exposure to environmental carcinogenic factors (*3, 4*). Only a smaller proportion of all cancer cases are associated with inherited genetic determinants (hereditary cancer), while their majority (sporadic cancer) is linked to environmental factors, which remain partially understood (*5, 6*). Environmental carcinogens are difficult to identify, and exposure to already known carcinogens is often not confirmed for individual cancer patients (*7*), suggesting significant gaps in our understanding of cancer initiation. Most known carcinogenic substances are genotoxic. Genotoxic chemicals induce DNA damage, resulting in mutations in somatic cells, the accumulation of which can fuel malignant transformation. In recent years, it has been shown that accumulation of somatic mutations can even be induced by the activation of endogenous Long Interspersed Element-1 (LINE-1 or L1) retrotransposons (*8, 9*). In this context, an important question is to what extent the mutagenic L1 activity occurs in somatic tissues and what effects might increase its intensity.

L1 retrotransposons are the only currently active autonomous mobile genetic elements in humans, reaching more than 500,000 copies and occupying approximately 17% of the genome (*10, 11*). They are found in all placental mammals, in most of which they are still active today (*12*). In mice, for example, they approach 600,000 copies and occupy about 19% of the genome (*13*). L1 elements form new genomic insertions by a “copy and paste” mechanism, known as retrotransposition, involving both encoded proteins (L1-ORF1p and L1-ORF2p) and occurring via an RNA intermediate (*10, 14*). L1 retrotransposition activity can alter the genomic structure in myriad ways. For example, a new L1 insertion may directly influence the transcript levels of nearby genes by promoter addition or disruption/introduction of cis-regulatory elements (*15*). Therefore, new L1 integrations often have a greater impact on the function of nearby genes than a point mutation.

For a long time, L1 elements were thought to be active only in the germline and early embryo. Such germline activity accelerates the evolution of mammalian genomes (*15-17*). In return, the new germline L1 integrations may result in inherited monogenic disorders in an individual (*18*). The biological significance of early embryonic L1 activity inducing somatic mosaicism is currently unknown (*19*). L1 retrotransposition activity is thought to be abolished later in embryonic development and remains undetectable in most normal somatic tissues due to several partially revealed cellular mechanisms that protect against somatic L1 activity (*20-28*). An exception is brain tissue, where L1 retrotransposition is detectable (*29*) and potentially associated to neuronal progenitor cells (*30*). Recently, sporadic L1 mRNA expression has also been detected in some epithelial cells (*31*). However, it is not known whether this leads to retrotransposition. L1 mRNA expression is likely to lead at least to translation of L1 proteins in epithelial cells, as similar sporadic L1-ORF1p immunopositivity can also be seen for example in normal cervical epithelium (*32*). The number of cells that allow L1 expression in normal tissues might increase with age. It was shown that, L1 mRNA expression is upregulated and the frequency of L1-ORF1p immunopositive cells is increased in tissues of aged mice (*33*). However, whether and to what extent L1 expression in normal young and old somatic tissues leads to L1 retrotransposition remains an open question, as there are several post-translational mechanisms of L1 regulation that may yet be able to prevent this (*24-27*). In sharp contrast with the normal somatic tissues, both marked L1 expression and retrotransposition are present in more than 50% of cancers (*9, 34*). In line with this, several cancer driver mutations have been identified in different tumor types that were caused by novel somatic L1 integration events (*9, 35-40*). Notably, L1 expression, and even retrotransposition in some studies, was shown in various cancer precursor syndromes (*32, 36, 41, 42*), suggesting an early cancer initiation role for L1 activity.

Currently, it is not clear how much basal L1 activity is present in somatic tissues and what might elevate it to higher intensities, as is already evident in many cancer precursor lesions. It would therefore be important to accurately measure the extent of somatic L1 retrotransposition. In principle, ORFeus-type L1 reporters (*43, 44*) appeared to be suitable for this measurement, but their application faced serious difficulties in detecting somatic L1 activity. These difficulties arise mainly from the fact that previous approaches have relied on the use of the ORFeus reporter in germline transgenic mice. In this setting the reporter gets activated during the period of physiological germline and early embryonic L1 retrotranspositions, long before somatic tissue development (*19*). Which, due to the lineage tracing nature of the signal produced by the activated ORFeus reporter, strongly interferes with the detection of later adult somatic L1 activity. However, studies using tet-inducible ORFeus reporters indicate that ORFeus activity is readily detectable when induction occurs during embryonic development, whereas substantially lower levels are observed upon induction at birth, implying strong negative regulation of retrotransposition in the soma that develops very early in life (*28*).

We have attempted to partly answer the remaining questions about L1 retrotransposition in normal somatic tissues by generating an assay system that avoids the pitfalls encountered with previous experimental approaches. Earlier, we developed a liver-specific somatic transgenesis procedure that enables stable transgene expression in the entire hepatocyte population of the murine liver throughout the lifespan of experimental animals (*45*). The stable and organ-wide transgene expression is facilitated by liver-targeted fumarylacetoacetate hydrolase (*Fah*) gene replacement induced liver regeneration in *Fah* knockout (KO) mice. The procedure also allows linkage of the expression of any transgene of interest to the expression of the *Fah* selection marker using the bidirectional HADHA/B promoter (*45*). We have used this procedure to achieve organ-wide somatic expression of ORFeus-type L1 reporters, avoiding the need for them to traverse the germline. This approach allows for the exclusive measurement of somatic L1 activity, separating it from the activity observed in germ cells and early embryos. We have applied modified ORFeus-type reporter elements, which support immunohistochemical (IHC) detection of cells harbouring reporter retrotransposition events. To better quantify the IHC-based detection of reporter retrotransposition, we have developed a novel artificial intelligence (AI)-based image analysis workflow. In addition to using traditional autonomous ORFeus reporters, we have also created non-autonomous (NA) reporter variants. These can be used to rule out the possibility that any somatic retrotransposition observed is solely due to the forced expression of autonomous reporters. Our results indicate that a basal L1 retrotransposition activity exists in normal organs, the intensity of which is presumably enhanced in response to various stimuli. Our novel assay could in the future be utilized for the identification of carcinogenic effects exerted through triggering or enhancing somatic L1 activity.

## Results

### Creation of novel ORFeus reporter variants and their characterization in mammalian tissue culture

The *in vivo* investigation of L1 reporter activation in animal tissues, down to the level of individual cells, would be most feasible by IHC-based detection of the signal generated by reporter retrotransposition. ORFeus-type reporters that produce an EGFP signal upon activation, such as the L1 reporter in the pWA125 vector (*46*)–a derivative of the native mouse L1spa element (*47*)–may be ideal for this purpose. However, a closer examination of the structure of the EGFP reporter cassette (*48*) in the traditional ORFeus element predicts that the large non-fluorescent EGFP fragment encoded by the first EGFP exon will be co-expressed in cells that carry the reporter, regardless of whether they have undergone reporter retrotransposition (Supplementary material 1 and SFig. 1). We confirmed the presence of such a transcript capable of expressing 3’-truncated EGFP in cultured HT1080 cells using RT-qPCR (Supplementary material 1 and SFig. 1). This poses a risk of unwanted background appearance during EGFP IHC staining in *in vivo* L1 reporter applications. To rule out this possibility, we developed a novel modified variant (ORFeus-IS) of the original mouse ORFeus element armed with an EGFP-based retrotransposition indicator cassette (*43, 46*). In the new ORFeus-IS the position of the intron separating the exons encoding the EGFP protein was shifted upstream within the coding sequence, resulting in a markedly smaller EGFP exon 1 (Fig. 1A). Consequently, the smaller co-expressed 3’-truncated EGFP fragment no longer displays the epitopes recognized by most EGFP antibodies (Supplementary material 1 and SFig. 3). This minimal modification eliminates the possibility for the formation of unwanted EGFP IHC background staining during *in vivo* applications. Both the original and the ORFeus-IS reporters were used to generate their non-autonomous (NA) variants (Fig. 1A and SFig. 3). In these variants, the reporters do not encode the L1-ORF2p essential for their retrotransposition. Successful retrotransposition of these variants requires the expression of the L1-ORF2p from endogenous L1 loci (SFig. 3).

**Fig. 1.**
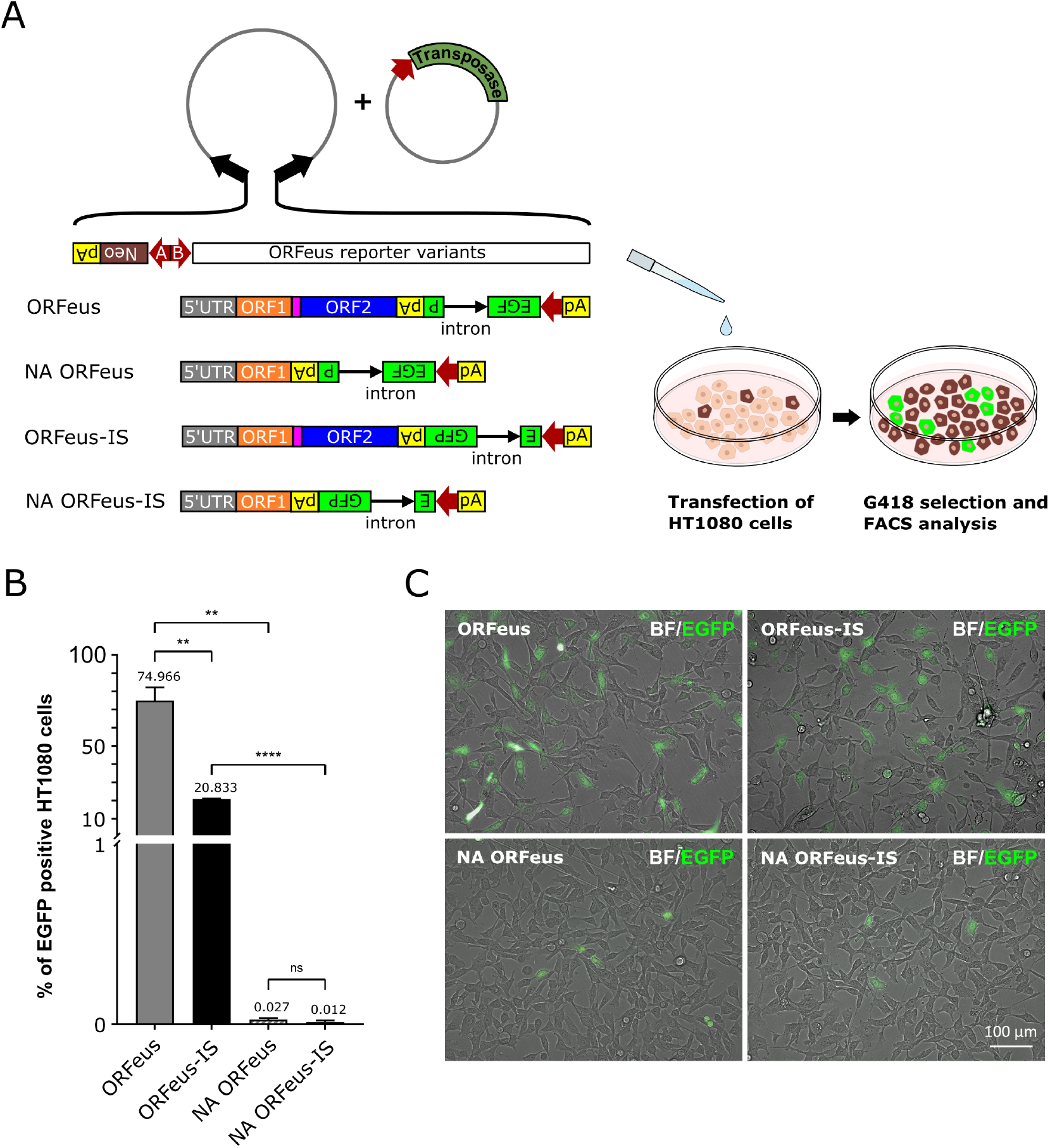
*In vitro* characterization of the traditional and novel ORFeus variants in mammalian cell culture. **(A)** Schematic representation of the structure of the *Sleeping Beauty* (SB) transposon-based cloning platform and outline of the mammalian cell culture experiments. Black arrows, SB transposon inverted terminal repeats; red arrows, promoters; pink rectangle, inter-ORF spacer. **(B)** Flow cytometry measurement of EGFP fluorescence in HT1080 cells stably transfected with the traditional (ORFeus) and the three new (NA ORFeus, ORFeus-IS, and NA ORFeus-IS) reporter variants. Data were presented as the mean ± standard deviation (SD) (n = 3) (see Supplementary Materials 2 for individual data values and statistics). **(C)** Representative overlay images of HT1080 cells stably transfected with all four L1 reporter variants imaged in brightfield and for green fluorescence.

All four L1 reporter variants were transfected into HT1080 cells to characterize and compare their functionality. The expression of ORFeus reporter variants was linked to that of the Neo selection marker gene using the bidirectional HADHA/B promoter (*45*) (Fig. 1A). For stable transfection and chromosomal gene transfer all four ORFeus variants were cloned between *Sleeping Beauty* (SB) transposon Inverted Terminal Repeats (ITRs) and the SB100 transposase helper (*49*) was added in trans (Fig. 1A). The extent of retrotransposition activity of the reporter is proportional to the number of EGFP positive cells, since full-length fluorescent EGFP protein is only produced after successful ORFeus retrotransposition (SFig. 1 and 3). After 1 month of G418 selection we measured the proportion of cells expressing EGFP by flow cytometry (Fig. 1B). The activity of NA variants was significantly lower—by more than three orders of magnitude—in HT1080 cells compared to that of their autonomous counterparts. (Fig. 1B). The activity of the ORFeus-IS elements decreased slightly compared to the original ORFeus reporters, producing retrotransposition over the same incubation time in 3.6 times and 2.2 times less HT1080 cells for the autonomous and non-autonomous variants, respectively (Fig. 1B). The retrotransposition activity of the different reporter variants was verified by microscopy in all cases (Fig. 1C). In conclusion, both the new ORFeus-IS and NA ORFeus-IS variants are active in HT1080 cells, where they show activity similar to their original ORFeus and NA ORFeus counterparts. Thus, they can be used to study L1 retrotransposition *in vivo*. In the next step, we tested the ORFeus-IS and NA-ORFeus-IS novel reporter variants in a somatically transgenic animal model.

### Somatic application of the novel ORFeus-IS reporter variants in mice

For single-cell-level examination of ORFeus-type reporter activation in animal tissues, we used the new ORFeus-IS and NA-ORFeus-IS reporter variants. Earlier, we developed a liver-specific somatic transgenesis procedure enabling stable transgene expression in the entire hepatocyte population of experimental mice throughout their lifespan (*45*). We have used this procedure to achieve organ-wide expression of the reporters avoiding the need for them to traverse the germline. Effective chromosomal gene transfer is essential for the somatic transgenesis protocol (*45*). We compensated for the reduced gene transfer efficiency caused by the large size of the ORFeus transgenes by switching to the *PiggyBac* (PB) transposon system, which is less sensitive for transgene size as compared to SB (*50, 51*). Similar to the ones used in mammalian tissue culture experiments, the central element of the reporter constructs used in *in vivo* testing was also the HADHA/B bidirectional promoter. However instead of Neo, we inserted a bicistronic unit expressing mCherry and Fah proteins on its A side (*45*) (Fig. 2A). On the B side of the promoter, we built the ORFeus-IS or the NA-ORFeus-IS reporter variants and placed the entire assembly between PB transposon ITRs (Fig. 2A). To test this arrangement *in vivo*, the L1 reporter constructs and a hyPB transposase helper (*52*) expressing plasmid were co-delivered hydrodynamically into the liver of female Fah deficient (*Fah*^*−*^*/*^*−*^) mice, then the curative drug nitisinone (NTBC) (*53*) was withdrawn to initiate liver regeneration (Fig. 2A). The *Fah*^*−*^*/*^*−*^ genetic background of the recipient animals imposes a strong selective pressure following NTBC withdrawal, favouring hepatocytes in which Fah function has been restored. Liver regeneration is initiated from these Fah-corrected hepatocytes. During the resulting multinodular regenerative process, Fah-deficient hepatocytes undergo cell death in the absence of NTBC, thereby being progressively replaced by proliferating transgenic hepatocytes. As a consequence, within slightly more than two months, a liver composed almost entirely of transgenic hepatocytes is generated. In this system, sustained expression of the transgenes is ensured by positive selection, facilitated by the use of the bidirectional promoter (*45*).

**Fig. 2.**
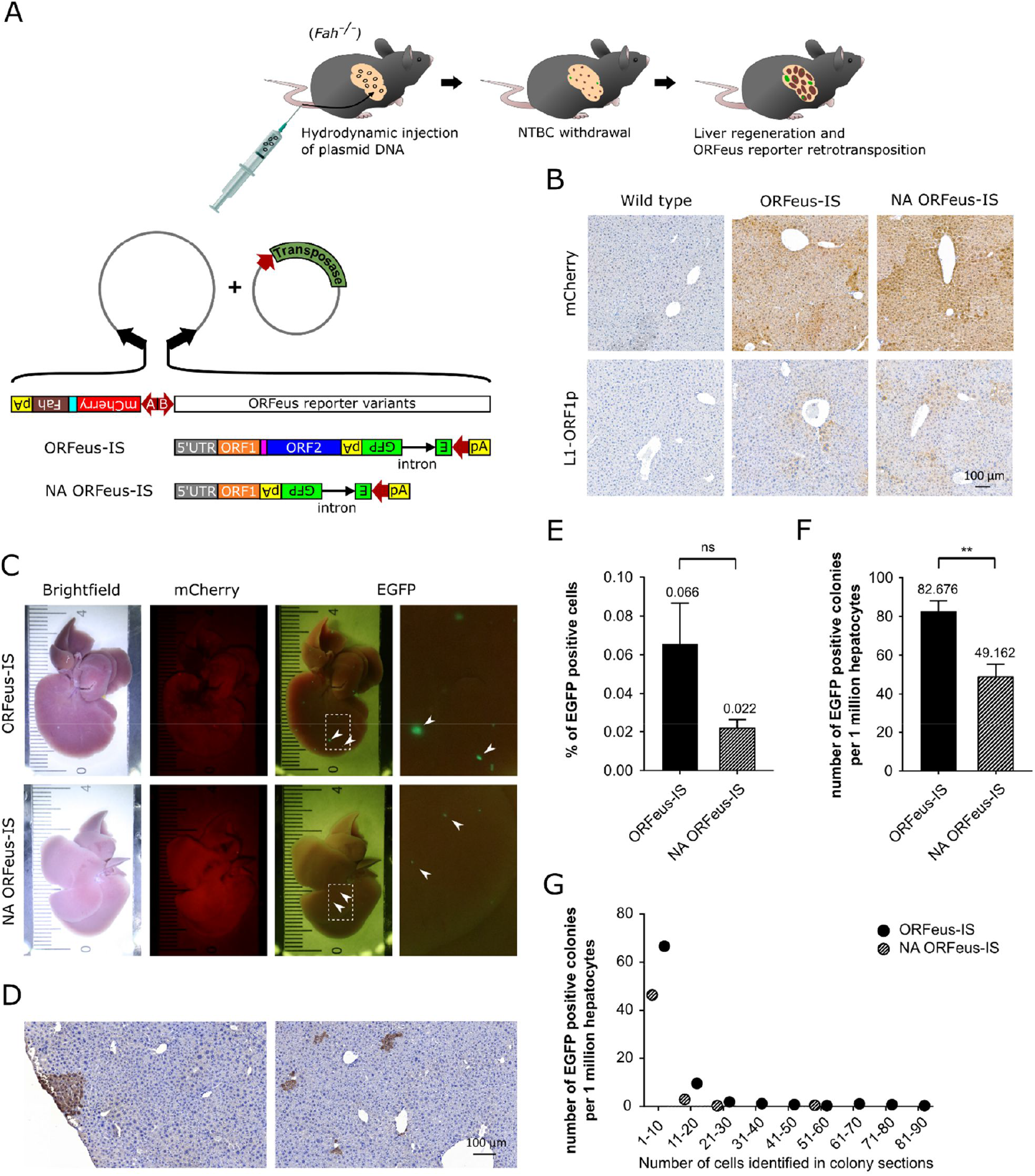
Assessment of the activity of the novel ORFeus-IS and NA ORFeus-IS reporter variants in the mouse liver. **(A)** Schematic representation of the structure of the *PiggyBac* (PB) transposon-based cloning platform and animal treatments. Black arrows, PB transposon inverted terminal repeats; red arrows, promoters; pink rectangle, inter-ORF spacer. **(B)** L1-ORF1p and mCherry immunostainings of liver sections from *Fah*^*−*^*/*^*−*^ mice 3 months after hydrodynamic injection of ORFeus-IS and NA ORFeus-IS elements and from wild type mice as controls. **(C)** Brightfield and fluorescence stereomicroscopic images of the liver of *Fah*^*−*^*/*^*−*^ mice 3 months after the intrahepatic delivery of ORFeus-IS and NA ORFeus-IS elements. **(D)** Representative EGFP immunostainings of liver sections from *Fah*^*−*^*/*^*−*^ mice 3 months after hydrodynamic injection of the ORFeus-IS element. **(E)** Determination of the percentage of EGFP-immunopositive hepatocytes carrying L1 reporter retrotransposition events using machine learning-based image analysis. EGFP immunostaining was performed on liver sections from *Fah*^*−*^*/*^*−*^ mice 3 months after hydrodynamic injection of ORFeus-IS and NA ORFeus-IS elements. Data were presented as the mean of 3 animals ± standard deviation (SD) (see Supplementary Materials 2 for individual data values and statistics). **(F)** Determination of the number of hepatocyte colonies consisting of EGFP-imunnopositive cells using machine learning-based image analysis. EGFP immunostaining was performed on liver sections from *Fah*^*−*^*/*^*−*^ mice 3 months after hydrodynamic injection of ORFeus-IS and NA ORFeus-IS elements. Data were normalized to 1 million cells and presented as the mean of 3 animals ± standard deviation (SD) (see Supplementary Materials 2 for individual data values and statistics). **(G)** Classification of the EGFP-immunopositive hepatocyte colonies based on their size. For each colony, we determined the number of associated EGFP-immunopositive hepatocytes in the examined section plane. Then colonies with different cell numbers were counted for all animals and colony counts were normalized to 1 million cells for each animal, taking into account the total number of hepatocytes examined for that animal. The number of colonies with a given cell count were presented in bins combining 10 cell count values as the mean of 3 animals (see Supplementary Materials 2 for individual data values and statistics).

Three months after hydrodynamic injection, following complete multi-nodular liver repopulation (*45, 54*), the animals were sacrificed. Sustained expression of the ORFeus-IS reporters in the mouse liver was indeed achieved as immunohistochemical (IHC) investigations demonstrated that virtually all hepatocytes were L1-ORF1p-immunopositive in the treated livers (Fig. 2B). Macrovisualization of mCherry fluorescence in the liver confirmed the success of organ-level transgenesis (*45*), while the patchy appearance of EGFP fluorescence indicated the somatic activation of L1 reporter elements (Fig. 2C). These results provide evidence that somatic L1 retrotransposition is detectable in the liver, even when a non-autonomous L1 reporter is used (Fig. 2C). The autonomous ORFeus-IS element produced more EGFP-immunopositive spots than the NA-ORFeus-IS (Fig. 2C) but the difference between the two constructs was remarkably smaller than that observed in the tissue culture context (Fig. 1B). Reporter retrotransposition was confirmed at the DNA level by an intron-spanning PCR assay that is able to detect loss of the EGFP intron in newly transposed L1 reporter copies (SFig. 4A). The assay confirmed the presence of newly transposed L1 reporter copies in the liver for both the ORFeus-IS and NA-ORFeus-IS variants (SFig. 4B).

In the next step, we set up the detection of full-length fluorescent EGFP by IHC, which allows for the quantitative identification of cells carrying ORFeus-IS or NA-ORFeus-IS reporter retrotransposition events. For ORFeus-IS elements, selective IHC-based detection of full-length EGFP, distinct from the polypeptide encoded by EGFP exon 1 (SFig. 3), was indeed possible (Fig. 2D). Although macro visualization of mCherry and EGFP using the original ORFeus elements yielded results similar to those produced with ORFeus-IS elements (compare SFig. 5B and Fig. 2C), we were unable to obtain valuable IHC outcomes with the original ORFeus elements (SFig. 5C). This further supports the presence of the reverse intron-retaining EGFP transcript (SFig. 1) and the interfering effect of the polypeptide encoded by the first EGFP exon in IHC studies when the traditional ORFeus elements are used.

Results of EGFP IHC examinations using ORFeus-IS elements further confirmed that somatic L1 retrotransposition is detectable in mouse liver (Fig. 2D). The number of EGFP-immunopositive cells was somewhat higher for the ORFeus-IS reporter than for NA-ORFeus-IS, consistent with the results of the EGFP fluorescence investigations (Fig. 2C). To accurately determine the number of liver cells carrying L1 reporter retrotransposition events, we evaluated the EGFP IHC pictures using machine learning-based image analysis. We found that when using the ORFeus-IS reporter, there was approximately 1 EGFP-immunopositive hepatocyte per 1700 hepatocytes (0.06%), while with NA-ORFeus-IS, there was approximately 1 per 5000 hepatocytes (0.02%) (Fig. 2E). This meant an approximately 3-fold difference between the two constructs in favour of the autonomous ORFeus-IS. This further strengthens the notion that the difference between the two constructs is much smaller in the liver than in cultured HT1080 cells (Fig. 1B). Our somatic transgenesis procedure coupled with liver regeneration also allows to analyse the number and size of hepatocyte colonies consisting of EGFP-imunnopositive cells (Fig. 2D). In this regard, we found a total of 1.7 times more colonies in the livers of mice expressing the ORFeus-IS element than in the livers of mice expressing NA ORFeus-IS (Fig. 2F). In a more detailed breakdown by size, the vast majority of colonies (more than 80%) contained 10 or fewer cells in the section plane examined (Fig. 2G and Supplementary material 2). Likewise, the proportion of colonies consisting of a single positive hepatocyte was also considerable (more than 25%) (Supplementary material 2).

## Discussion

The impact of L1 activity on the host organism is still far from fully understood. In this regard, it is important to distinguish between L1 activity in germline versus somatic tissues and the effects of L1 expression versus L1 retrotransposition. In general, L1 expression involved in stress responses can be a mediator of both beneficial and harmful cellular processes. Its potential beneficial effect is exemplified by its recently discovered positive role in osteogenic repair (*55*). Similarly, in the context of cancer, it has been shown to trigger cell death in premalignant cells by activating the innate immune response (*56*). At the same time, L1 expression can also play a role in harmful processes like autoimmune disorders (*57, 58*) and aging-associated inflammation (*33*). In cancer, massive L1 expression has been described in several cancer precursor syndromes (*32, 36, 41*), where this expression did not lead to the elimination of premalignant cells and persisted in mature tumors during the course of carcinogenesis. Conversely, L1 retrotransposition is more clearly harmful, as its genome-destabilizing effect promotes tumorigenesis in somatic tissues by creating new L1 insertions and inducing other types of mutations (*9, 59*). Therefore, it is important to study the dynamics of L1 retrotransposition in somatic tissues.

In this study, we examined L1 retrotransposition activity using autonomous and non-autonomous L1 reporter variants *in vitro* in cultured human cells and *in vivo* using a somatic gene delivery system in mice designed for this purpose. The results obtained in mouse somatic tissue model are also relevant in humans, since basic features of L1 biology are similar in mice and humans and many somatic L1 control mechanisms are also known to be shared between the two species (*20-22, 24, 60*). L1 reporter constructs used for either *in vitro* pre-testing or *in vivo* study were designed to enable both DNA transposon-based stable gene delivery and generation of positive selection pressure for L1 reporter expression utilizing a bidirectional promoter (*45*). Theoretically, such forced expression of ORFeus reporters may allow the artificial induction of L1 reporter retrotransposition events even in the presence of fully functional cellular mechanisms protecting against L1 activity. To control for this, we decided not to express the entire ORFeus element, but rather a variant (NA) that only encodes L1-ORF1p, thereby rendering it non-functional on its own due to the absence of L1-ORF2p expression (SFig. 3). The expression of L1-ORF2p, necessary for the function of these NA ORFeus reporter variants, is under physiological epigenetic control in endogenous loci, and is free from the positive selection pressure generated by our constructs. According to our results, ORFeus reporters carrying the original EGFP reporter cassette (*46*) cannot be reliably used for IHC-based evaluation in *in vivo* studies, as their use may lead to unwanted IHC background staining (SFig. 5C). We solved this problem by slightly modifying the ORFeus element, solely in the structure of the EGFP reporter cassette. In the new ORFeus-IS variants, the position of the intron in the EGFP coding sequence was shifted upstream (SFig. 2 and 3), which enabled stable IHC-based evaluation (Fig. 2D and SFig. 5C).

We pre-tested the new ORFeus reporter variants in the HT1080 human fibrosarcoma cell line. Mouse L1 elements function in human cells and vice versa, without a major loss of activity (*61, 62*). Moreover, the human and mouse L1-ORF1p and L1-ORF2p proteins are interchangeable. The retrotransposition activity of human-mouse L1 element chimaeras generated by the interspecies exchange of L1-ORF1p or L1-ORF2p is not significantly reduced in either host (*63*). Thus, measurements performed on human cell lines are relevant to mice and vice versa for all tested reporter variants. Using the original ORFeus reporter element, after 1 month of culturing under continuous G418 selection, we found 75% EGFP-positive HT1080 cells carrying L1 reporter retrotransposition events (Fig. 1B). This high value is roughly consistent with the experiences of others using similar L1 reporter elements in combination with various gene delivery systems in cultured cells (*64, 65*). In contrast, for the NA ORFeus variant that does not express L1-ORF2p, we measured a value of 0.03%, representing a strong 3-order of magnitude decrease in retrotransposition activity (Fig. 1B). The same difference was observed between the ORfeus-IS and NA ORFeus-IS variants (Fig. 1B). The drop in the activity of the NA variants compared to that of the parental ORFeus and ORFeus-IS elements could be explained by the fact that their retrotransposition requires the expression of L1-ORF2p transcribed from other endogenous loci and its entry into the reporter mRNA/L1-ORF1p RNP complex (SFig. 3).

During the *in vivo* study, we intended to avoid the interfering effect of the physiological germline and early embryonic L1 activity (*19, 66*). This prior activity, together with the lineage tracing nature of the ORFeus reporters, makes it problematic to monitor somatic L1 activity in germline-transgenic mouse models. Therefore, we used a somatic gene delivery technology enabling long-lasting transgene expression in the entire hepatocyte population of mice (*45*). The technology induces multi-nodular liver repopulation from transgenic cells, which is completed approximately 2 months after gene delivery, creating a transgenic liver in its entire hepatocyte population (*45, 54*). With this technology, we could detect somatic L1 retrotransposition in the mouse liver, even when using the NA ORFeus-IS element (Fig. 2C-E). The results were obtained at the HUN-REN BRC, Central Animal House facility under the current standard animal husbandry conditions. Thus, since the chemical factors affecting somatic retrotransposition are not yet known, deviation from this is possible under other animal husbandry conditions. Our results suggest that somatic blockade of L1 retrotransposition is leaky, and thus genome-destabilizing *de novo* L1 retrotranspositions occur in somatic tissues. Another recent publication also refers to retrotransposition occurring in normal human colorectal epithelium (*67*).

Surprisingly, the proportion of EGFP-positive cells obtained in the mouse liver (Fig. 2E) was in the same range for both ORFeus-IS (0.066%) and NA ORFeus-IS (0.022%) elements as for NA ORFeus-IS (0.012%) elements in the tumor-derived cell culture (Fig. 1B). From this activity range, only the results obtained with the ORFeus-IS (20.833%) element in cancer cell culture stand out by 3 orders of magnitude (Fig. 1B). The comparability of tissue culture measurements and *in vivo* investigations is given by the fact that in both cases the same reporter variants were examined with the same construct structure, where we placed positive selection pressure on L1 reporter expression (Fig. 1A and Fig. 2A). In addition, during the 1-month G418 selection, the cell cultures examined underwent a similar number of cell division as transgenic hepatocytes during liver regeneration (*45, 53*). The high ORFeus-IS activity measured in HT1080 cells is consistent with the experience of other laboratories using similar ORFeus-type reporters in cultured cells (*64, 65*). The other three measurements performed by using NA reporter variants and *in vivo* in the mouse liver are unprecedented in the scientific literature (Fig. 3). We obtained similar results with NA reporter variants in other human and mouse tumor-derived cell lines (unpublished data). Considering that L1-ORF2p expression is under endogenous epigenetic control in the case of NA-type reporters, these findings suggests that tumor cells primarily rely on epigenetic regulation to restrict L1 retrotransposition, while downstream defense mechanisms are more permissive, likely due to their impaired function (Fig. 3). In contrast, in normal liver tissue, ORFeus-IS activity remained in the low range typical of NA ORFeus-IS elements, showing only a 3-fold increase compared to the activity of its NA counterpart. These observation implies that, in normal somatic tissues, tight epigenetic control is not the primary barrier to L1 retrotransposition. Instead, downstream regulatory mechanisms acting after L1 expression appear to function more efficiently, thereby limiting productive retrotransposition events (Fig. 3). This model is consistent with the large number of endogenous L1 loci present in mammalian genomes and with reports showing that L1 expression is frequently induced during stress responses in somatic tissues (*55, 57*). Together, these findings indicate that L1 regulation in tumor-derived cell cultures operates under fundamentally different constraints than in normal somatic tissues.

**Fig. 3.**
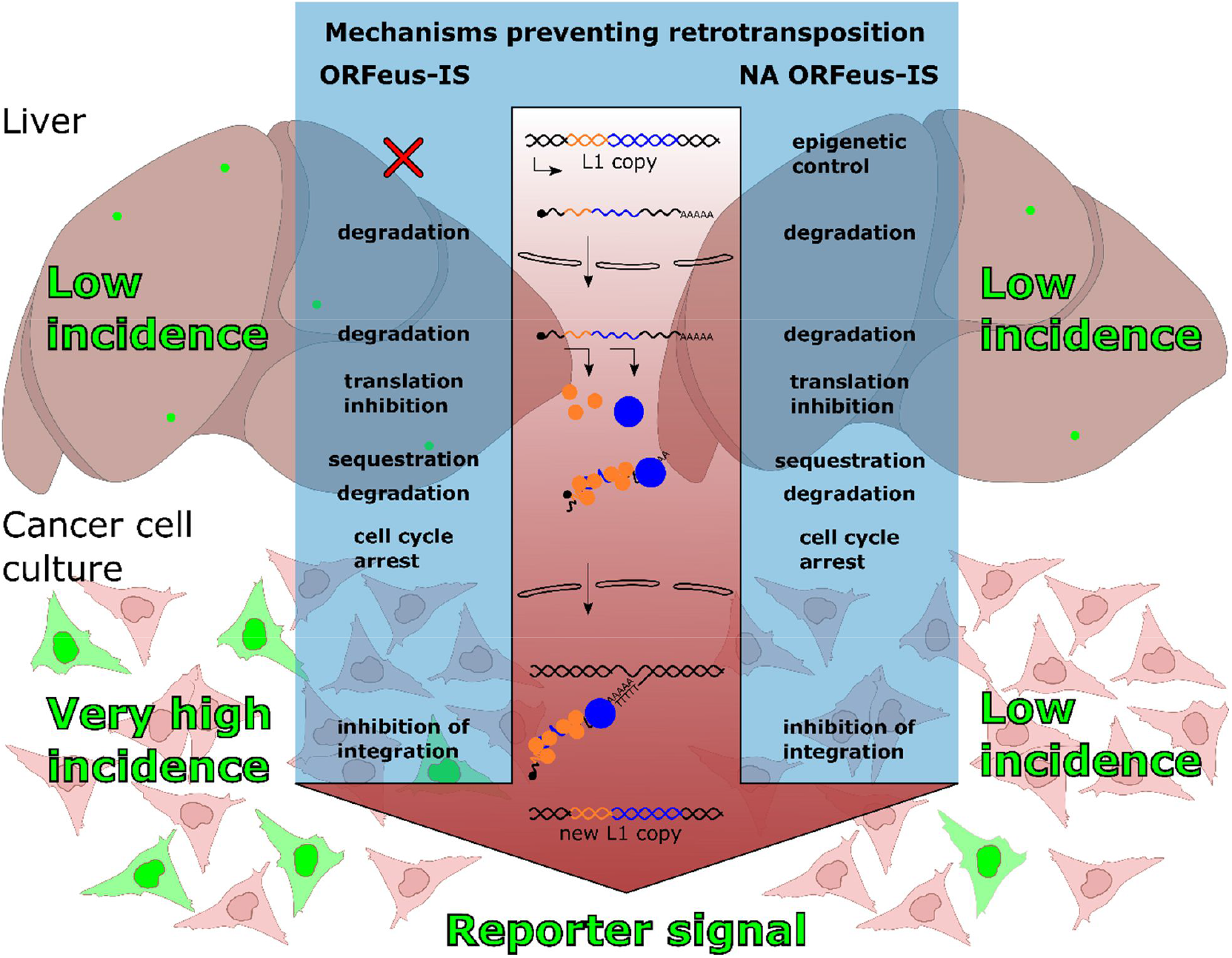
Schematic representation of the L1 retrotransposition mechanism and somatic control. L1 retrotransposition and the mechanisms controlling it are shown in light of the use of autonomous and non-autonomous L1 reporters, providing background for interpreting reporter activity results obtained in *in vitro* and *in vivo* gene delivery experiments. ORFeus-IS reporter activity represents exogenous L1 activity, while NA ORFeus-IS reporter activity represents endogenous L1 activity, since in the latter case, L1-ORF2p expression is under endogenous epigenetic control.

In the liver we also analyzed the number and size of colonies formed by EGFP-positive hepatocytes (Fig. 2F and G). IHC images show colonies of varying sizes, ranging from those containing around 100 cells down to single cells in the section plane examined (Fig. 2D and Supplementary material 2). In tissue culture experiments, while a similar number of cell divisions occur during 1 month of culturing, the cells are replated several times, separating the descendants from each other, whereas during liver regeneration they remain adjacent and form colonies. The size of the colonies indicates the time of the L1 reporter retrotransposition event during the course of liver regeneration. Large colonies form as a result of early events, while single cells are late events, potentially occurring after regenerative cell divisions have been completed. The size distribution of colonies obtained using ORFeus-IS and NA ORFeus-IS reporters was comparable (Fig. 2G). There was a large number of both identified single EGFP-positive cells and colonies with 4 or fewer cells (Supplementary material 2). The presence of such a large number of individual cells suggests post-regenerative L1 reporter retrotransposition events. They could potentially form because the animals were sacrificed weeks after the completion of liver regeneration, 3 months after gene delivery. The higher proportion of colonies with low cell numbers can probably be attributed to multiple reasons. As cell divisions progress, the number of hepatocytes carrying the L1 reporter increases exponentially, enabling retrotranspositions to occur in more cells at later stages of regeneration. In addition, the rate of cell division slows down in the later stages of liver regeneration (*68*), thus allowing more time for reporter retrotranspositions to occur than during early regenerative divisions. Furthermore, it is also known that even in non-regenerating normal liver, liver homeostasis is maintained by modest hepatocyte proliferation (*69*).

Our results suggest that L1 retrotransposition might be more common in mammalian somatic tissues than previously thought. In principle, a variety of factors, such as infections, chemicals, etc., can disturb some of the mechanisms that protect against L1 activity (Fig. 3) The resulting increase in L1 retrotransposition burden may generate driver mutations with increased probability in somatic tissues to fuel the process of carcinogenesis. Our intrahepatic L1 reporter assay could be used in the future to identify such factors, leading to earlier hazard identification and better cancer prevention on a population level.

## Materials and Methods

### Animal care and maintenance

Mice were bred and maintained in the Central Animal House at the HUN-REN Biological Research Centre (Szeged, Hungary). The specific pathogen-free status was confirmed quarterly according to FELASA (Federation for Laboratory Animal Science Associations) recommendations. All animal experiments were conducted in accordance with the guidelines of the Institutional Animal Care and Use Committee at the HUN-REN Biological Research Centre under the supervision of the Governmental Office for Csongrád-Csanád County, Directorate of Food Chain Safety and Animal Health. The approval number is XVI./2396/2023.

### Plasmid construction

The transposase helper expressing pcGlobin2-SB100 and pcGlobin2-hyPB plasmids were constructed as described (*49, 54*). For the creation of the ORFeus-IS and NA ORFeus L1 reporter variants the MCS of the pUC57-simple plasmid backbone was replaced by NotI and SgfI REN recognition sites, then the 10188 bp NotI-PvuI fragment containing the complete ORFeus reporter from the pWA125 (*46*) plasmid was cloned into this plasmid backbone resulting in pUC57-ORFeus125. To create ORFeus-IS reporter variants, the 2333 bp AgeI-NheI fragment containing the hsvTK polyadenylation signal, the EGFP exons and the modified human γ-globin intron of the retrotransposition indicator cassette were synthesized by GenScript (GenScript, Piscataway, NJ, USA) with the modification that the intron position was shifted 375 bp (125 AA) upstream within the EGFP coding sequence. The 2333 bp AgeI-NheI fragment was then replaced with the resynthesized fragment in the pUC57-ORFeus125 plasmid. To create NA ORFeus reporter variants, the 3874 bp PacI-AgeI fragment containing the L1-ORF2p coding sequence was deleted in the pUC57-ORFeus125 plasmid. The final vector variants used for testing were designed to enable both DNA transposon-based stable gene delivery and generation of positive selection pressure for L1 reporter expression utilizing a bidirectional promoter. To create L1 reporter variants for *in vitro* testing in mammalian cell culture, the promoter of the pNeo-miR vector (*45*) was replaced with the bidirectional HADHA/B promoter, so that its A side expressed the Neo transgene, while NotI and SgfI REN cleavage sites suitable for cloning the ORFeus elements were built next to its B side. Then, the NotI-PvuI fragment containing the complete ORFeus reporter element was cloned from the pUC57-ORFeus125 plasmid into NotI-SgfI sites on the B side of the bidirectional promoter to create the pUC57-SB-Neo-ORFeus construct. The pUC57-SB-Neo-NA ORFeus, pUC57-SB-Neo-ORFeus-IS and pUC57-SB-Neo-NA ORFeus-IS variants were created accordingly, using the modified pUC57-ORFeus125 vector variants. To create L1 reporter variants for *in vivo* testing in the mouse liver, the 3518 bp EcoRI-NotI fragment from the pbiLiv-miR vector (*45*) containing the HADHA/B promoter, the mCherry-T2A-Fah coding sequence with a modified EEF1A1 intron and a bGH polyadenylation signal was cloned between PB ITR elements in a way that the NotI and SgfI REN recognition sites suitable for cloning the ORFeus elements were placed on the B side of the HADHA/B promoter. Then, the NotI-PvuI fragment containing the complete ORFeus reporter element was cloned from the pUC57-ORFeus125 plasmid into NotI-SgfI sites on the B side of the bidirectional promoter to create the pUC57-PB-Fah-ORFeus construct. The pUC57-PB-Fah-NA ORFeus, pUC57-PB-Fah-ORFeus-IS and pUC57-PB-Fah-NA ORFeus-IS variants were created accordingly, using the modified pUC57-ORFeus125 vector variants.

### Cell culture and transfection

HT1080 cells were purchased from ATCC and cultured in Dulbecco’s modified Eagle’s medium (DMEM) (Biosera) supplemented with 10% fetal bovine serum (FBS; Gibco) and 1% penicillin-streptomycin (P-S; HyClone) in the presence of 5% CO2 at 37 °C. Cells were transfected with 500 ng of each reporter construct, in combination with 50 ng of the transposase helper plasmid, using FuGENE® HD transfection reagent (Promega) according to the manufacturer’s instructions. Selection of stable transfected cells was performed using neomycin (G418; Biosera).

### Flow cytometry

Neomycin (G418; Biosera) selection was performed continuously on the L1 reporter transfected HT1080 cells for 1 month. Then cells were trypsinised, pelleted (300×g, 5 min), and resuspended in 500 µL DMEM. Propidium iodide (PI, 1 µg/mL working concentration) was added, and samples were incubated for 5 min at room temperature, protected from light, to label necrotic cells with compromised membrane integrity before acquisition (*70, 71*). Cells were measured on a CytoFLEX S instrument (Beckman Coulter), and 50.000 singlet events per sample were collected. Data were gated for debris (FSC/SSC), singlets, and live cells (PI negative). EGFP-positive percentage was quantified within the live gate. Analysis was carried out with CytExpert (v.2.3, Beckman Coulter) analysis software.

### Hydrodynamic tail vein injection

Plasmids for hydrodynamic tail vein injection were prepared using the NucleoBond Xtra Maxi Plus EF Kit (Macherey-Nagel) according to the manufacturer’s instructions. Before injection, we diluted plasmid DNA in Ringer’s solution (0.9% NaCl, 0.03% KCl, 0.016% CaCl2) and a volume equivalent to 10% of mouse body weight was administered via the lateral tail vein in 5–8 s into 6– 8-week-old *Fah*^*−*^*/*^*−*^ (C57BL/6N-*Fahtm1(NCOM)Mfgc/Biat*) mice. The amount of plasmid DNA was 50 μg for each of the constructs mixed with 4 μg of the transposase helper plasmid. *Fah*^*−*^*/*^*−*^ mice were treated with 8 mg/L Nitisinone (NTBC) (Merck) in drinking water. After hydrodynamic injection with constructs expressing the Fah protein, NTBC was promptly withdrawn.

### RTqPCR experiments

Total RNA from cultured cells was isolated using TRI Reagent (MRC) following the manufacturer’s protocol. RNA was Dnase I-treated with PerfeCTa DNase I (Quantabio) and reverse transcribed into cDNA using RevertAid First Strand cDNA Synthesis Kit (ThermoFisher Scientific). RT-qPCR was performed on a Rotor-Gene Q instrument (Qiagen) with PerfeCTa SYBR Green Super Mix (Quantabio) as follows: 95 °C for 7 min followed by 40 cycles of 10 s at 95 °C, 15 s at 60 °C. All reactions were carried out in triplicate in a final volume of 20 μl. The primers EGFP intron fw (5’ AAGATACTGGGGTTGGGGT 3’) and EGFP intron rev (5’ CAGAGGCATTCCTTAGGCCC 3’) were used for the EGFP intron-specific assay and primers CMVE fw (5’ ACGCCAATAGGGACTTTCCA 3’) and CMVE rev (5’ TAGGGGGCGTACTTGGCATA 3’) were used for the CMV enhancer-specific assay. Detection of the ribosomal protein L27 (RPL27) mRNA served as a loading control using primers RPL27 fw (5’ CGCAAAGCTGTCATCGTG 3’) and RPL27 rev (5’ GTCACTTTGCGGGGGTAG 3’). PCR efficiencies were analysed with Rotor-Gene Q software (Qiagen). Gene expression was analysed by normalising to RPL27 using the ΔCT method (*72*).

### PCR-based detection of L1 reporter retrotransposition

Left liver lobes from reporter-bearing mice were lysed in 50 mL of lysis buffer (100 mM Tris-HCl, pH 8.0; 5 mM EDTA, pH 8.0; 200 mM NaCl; 0.2% SDS) and incubated overnight with 300 µg/mL proteinase K (VWR Chemicals). Genomic DNA (gDNA) was isolated from 1 mL of lysate by conventional phenol–chloroform extraction followed by ethanol precipitation. For detection of L1 retrotransposition, 1 µg of gDNA was used as template for standard PCR under the following cycling conditions: 98 °C for 30 s, followed by 35 cycles of 98 °C for 30 s, 67 °C for 20 s, and 72 °C for 15 s. The applied primers (EGFP fw: 5’ GAACTTGTGGCCGTTTACGTC 3’ and EGFP rev: 5’ CATGTGATCGCGCTTCTCGT 3’) hybridizing to the EGFP coding sequence, were designed to yield PCR products spanning the EGFP intron in the ORFeus-IS reporter variants. The PCR assay is producing a 1,496-bp intron-containing product over the original ORFeus-IS transgenes, and a 594-bp intronless amplicon over the secondary ORFeus-IS copies that have undergone retrotransposition. PCR products were resolved on a 1.5% agarose gel alongside a 1 kb DNA ladder (GeneRuler 1 kb, Thermo Fisher Scientific).

### Imaging

Pictures of whole mouse livers were taken with an Olympus SZX12 fluorescence stereozoom microscope equipped with a 100-W mercury lamp and filter sets for selective excitation and emission of GFP and mCherry. Representative images of HT1080 cells were acquired using an EVOS FLoid Cell Imaging System (Thermo Fisher Scientific) with brightfield and green channels. The original grayscale TIFF images for both channels were processed using GIMP (https://www.gimp.org/). Green-channel images were artificially colourized and overlaid with their corresponding brightfield images. Composite image layers were adjusted uniformly for brightness, contrast, exposure, shadows, and highlights.

### Immunohistochemistry

Mice were sacrificed 3 months after hydrodynamic injection. Livers were removed and fixed in 4% formalin overnight, dehidrated in an automatic benchtop tissue processor (Leica TP1020) and paraffinated (Leica, HistoCore Arcadia H). From fixed tissues, 5 μm sections were cut and incubated at 62 °C for 1 hour, prior to the staining. Deparaffinization was done in a Leica Autostainer XL machine. Antigen retrieval was done with Envision FLEX Mini Kit (DAKO) and Leica Novocastra Epitope Retrieval solutions. The primary antibodies used were rabbit polyclonal anti-EGFP (Abbexa, abx122699), preadsorbed on wildtype samples for 60 minutes and incubated on the samples for additional 30 minutes and rabbit monoclonal anti-LINE-1 ORF1p (Abcam, ab216324), incubated overnight on 4 °C. The secondary polyclonal rabbit anti-mouse-HRP (Leica, Novolink Polymer Detection Systems) antibody was incubated for 30 minutes. Visualization was done with EnVision FLEX DAB+ Chromogen System (DAKO, GV825). After hematoxylin counterstaining (Leica, Novolink Polymer Detection Systems) for 3 minutes, slides were mounted and scanned with a Pannoramic 1000 Digital Scanner (3DHistech).

### Image analysis pipeline

Slides were digitized using a Pannoramic 1000 Digital Scanner (3DHistech) at ×20 optical magnification. These digitized images were processed using BIAS (BioImage Analysis Software) (*73*). Individual nuclei were detected using the Nucleaizer basic nucleus segmentation model (*74*) integrated into BIAS. In segmentation post-processing, one additional region was defined for each nucleus to gain information from cytoplasm. This cytoplasmic region, representing the entire cell, was generated by dilating the nuclear region up to a maximum radius of 10 μm, ensuring that adjacent cells did not overlap. Finally, standard features describing morphology, intensity, and texture were extracted from both nuclear and cytoplasmic regions from all channels for subsequent cell classification. Supervised machine learning was employed to predict four distinct cell types. These classes were manually selected based on their morphological characteristics. Cells with small and dark blue nuclei were considered as lymphocyte-like immune cells. Small segmented regions outside the tissue section were also classified as trash. Hepatocytes with evenly distributed brown chromogen signal (anti-EGFP staining) across the whole cells were labeled as EGFP-positive, whereas hepatocytes lacking chromogen staining were labeled as EGFP-negative. For the training set, we annotated approximately 350 cells for each class from different tissue sections. Support Vector Machine (SVM) was trained with a radial basis function kernel commonly used for the multi-class cell phenotype classification. After training the SVM model, model performance was assessed using 10-fold cross-validation to estimate classification accuracy. The trained model was then applied to classify all remaining cells on each liver section. 0.5 to 1 million hepatocytes were identified in multiple sections from each experimental animal for further analysis. The number of EGFP-positive and EGFP-negative hepatocytes and the number of colonies consisting of EGFP-positive hepatocytes were determined. Adjacent EGFP-positive cells with a shared cytoplasmic border were merged into a single region, which was considered to represent one colony.

### Data visualisation and statistics

GraphPad Prism 10 software (version 10.6.1 for Windows, GraphPad Software) was used for data visualization. To identify levels of statistical significance Welch’s t-test was applied. The threshold for significance was *P*<0.05. For details of individual statistical tests, see Supplementary material 2.

## Supporting information

Individual data values and statistics

## Acknowledgments

The authors acknowledge the provided support of the Central Animal House, FACS and Cellular Imaging Laboratory Core Facilities at HUN-REN BRC.

## Funding

The project was supported by the 2021-1.1.4-GYORSíTÓSÁV-2022-00008 and 2021-1.1.4-GYORSíTÓSÁV-2022-00018 projects of the National Research, Development and Innovation Fund, Hungary (IN), TKP2021-EGA09 no. TKP-31-8/PALY-2021 (PH), TKCS-2024/73 (PH), Horizon-BIALYMPH no. 2023-1.2.1-ERA_NET-2023-00002 (PH), Horizon-SYMMETRY no. 2019-2.1.7-ERA-NET-2022-00044 (PH), Horizon-SWEEPICS no. 101135053 (PH), Horizon 2020-Fair-CHARM no. 101016457 (PH), HAS-NAP3 no. NAP2022-I-6/2022 (PH), HUNRENTECH no. TECH-2024-34, OTKA-SNN_21 no. 139455/ARRS (PH), NKKP no. ADVANCED_24_151202 and a GenScript Life Science Research Grant 2024 (LM).

## Author contributions

LM conceived the study, designed the experiments. LM and AMN contributed to the project supervision and administration. LM contributed to the analysis and interpretation of data with help from JDB, WA and AMN. LM, JDB, WA, and AMN contributed to the writing and editing of the manuscript. PH, IN, JDB, WA and ZL. provided support of key infrastructure, reagents, and critical discussion of the manuscript. MR, RK, AMN, and GI performed the animal experiments. AMN, MR, OOV, RK, AGK, DSK, and KV participated in the construction and purification of plasmids. GI performed the fluorescent microscopy imaging. MR, carried out RT-qPCR experiments. OOV, RK, FS and KV performed the immunohistochemistry. EM performed the image analysis. MR, GI, KS and PK carried out the cell culture experiments. All authors read and approved the final manuscript.

## Competing interests

LM, RK, AMN, GI, AGK, PH and FS have submitted a PCT application related to the somatic application of the ORFeus reporters in mice. Jef Boeke is a Founder and Director of CDI Labs, Inc., a Founder of and consultant to Opentrons LabWorks/Neochromosome, Inc, a Founder of JATech, LLC, and serves or served on the Scientific Advisory Board of the following: CZ Biohub New York, LLC; Logomix, Inc.; Rome Therapeutics, Inc.; SeaHub, Seattle, WA; Tessera Therapeutics, Inc.; and the Wyss Institute. All other authors declare they have no competing interests.

## Data and materials availability

Use of the pWA125 plasmid is restricted to non-commercial purposes under an MTA issued by Johns Hopkins University School of Medicine.

