## Supplementary Text and Figs. S1 to S5 for "Intrahepatic reporter assay reveals leaky somatic blockade of L1 retrotransposition in mice"

May Raya *et al.*

**This PDF file includes:**

Supplementary Text
Figs. S1 to S5
References (75 to 77)

### Supplementary Text

<u>Extended technical description on the creation of ORFeus-IS reporter variants</u>

The structure of the traditional mouse ORFeus element on the pWA125 plasmid vector (*45*) predicts that the large polypeptide encoded by EGFP exon 1 will also be present in all cells expressing the reporter, regardless of whether they have undergone ORFeus reporter retrotransposition (SFig. 1A). The ORFeus element contains the L1 5'UTR and the codon-optimized ORF1 and ORF2 sequences from the endogenous mouse L1spa (*46*) element and a sense intron-disrupted antisense oriented EGFP retrotransposition indicator cassette (*47*). The EGFP cassette is flanked by a CMV immediate early promoter and a herpes simplex virus thymidine kinase (hsvTK) polyadenylation signal on the antisense strand.

Theoretically, there is no obstacle preventing the transcription of the antisense EGFP reporter gene in an overlapping manner with the sense L1 transcription even in the case of the original ORFeus transgenes still containing the intron (SFig. 1A). Such overlapping sense-antisense (S-AS) transcript pairs are very common in mammalian genomes (*75*). The antisense EGFP reporter transcription can express the polypeptide encoded by EGFP exon 1 even from non-transposed ORFeus copies, extended by a short peptide added from the sense-oriented intron, up to the first stop codon (SFig. 1A). Although this does not produce fluorescent EGFP, it represents a theoretical possibility for the appearance of a background staining during EGFP immunohistochemistry (IHC). Such IHC background staining would complicate or prevent *in silico* image analysis for the quantitative processing of somatic ORFeus reporter activity.

In the antisense reporter transcript expressed by the EGFP cassette (*47*) (SFig. 1A), exon 1 encodes the first 178 amino acids (AAs) of EGFP (239 AAs in total), which is followed by a reverse-oriented (sense) modified human γ-globin intron that contributes 3 additional AAs (Leu, Leu, and Arg) before a TGA stop codon terminates translation (SFig. 2A). Since this 902 bp long intron cannot be recognized from the direction of the reporter transcript its retention results in a total length of 1243 bp sequence before reaching the hsvTK polyA addition signal (SFig. 1A and SFig. 2A). Such distance between the stop codon and the polyA signal does not suggest drastic transcript destabilization by nonsense-mediated mRNA decay (NMD) (*76*), thus it can be regarded as a transcript that displays the first three-quarters of the whole EGFP protein sequence. Although this 178 AA EGFP fragment encoded by exon 1 is not fluorescent, it carries the immunogenic epitopes recognized by most available EGFP antibodies (SFig. 1A). Consequently, its presence would hamper the distinctive IHC-based detection of the full-length EGFP protein appearing only following ORFeus reporter retrotransposition (SFig. 1A).

In order to enable stable chromosomal transgene delivery and positive selection for transgene expression, we cloned the mouse ORFeus element into *Sleeping Beauty* (SB) transposon (*47*) and linked its expression to that of the Neo positive selection marker, using the bidirectional HADHA/B promoter (*44*) (SFig. 1B). Upon transfection of this ORFeus reporter vector into HT1080 cells (SFig. 1B), following G418 selection we verified the presence of the reporter transcript retaining the reverse intron using an intron-specific RT-qPCR assay. The RT-qPCR experiments showed that the reporter transcript carrying the retained reverse intron is present in the ORFeus-expressing cells in approximately 70-fold lower amount than RPL27 (SFig. 1C). We also characterized the amount of the sense L1 transcript using a CMV enhancer region specific RTqPCR amplicon. This region is part of the L1 transcript but is missing from the reporter transcript. According to the results of these RTqPCR experiments, the amount of L1 transcript was approximately 160-fold lower than that of RPL27 (SFig. 1C). The amount of the intron excised from the L1 transcript is expected to be significantly less than that of the mature L1 transcript if the intron is only transcribed during the sense transcription. The rapidly degraded excised introns are generally detected in quantities that are on average one order of magnitude lower than those of their corresponding mature mRNAs (*77*). By contrast herein, the amount of intron-containing transcripts was more than twice that of the sense L1 transcript. This clearly indicates that the antisense reverse intron-retained reporter transcript is present in cells expressing the ORFeus reporter. It should be noted that using the intron specific RTqPCR assay endogenous γ-globin introns are also detectable in approximately 10,000-fold lower amount than RPL27 in HT1080 cells that do not express ORFeus element (SFig. 1C). The reason for this is that the intron-specific primers used, albeit with a few mismatches, also hybridize to the corresponding intron of the endogenous human γ-globin genes. However, this small quantity of endogenous transcripts detected here represent only a negligible fraction of the intron-specific RTqPCR values measured in ORFeus-transfected cells, thus not affecting our conclusions.

In the new ORFeus-IS reporter variant the position of the intron separating the exons encoding the EGFP protein was shifted 375 bp (125 AA) upstream within the EGFP coding sequence (SFig. 2B and SFig. 3). In the ORFeus-IS element, the hsvTK polyA signal is reached 1618 bp after the new, shorter EGFP exon 1. This modification ensures that a transcript containing only a 159 bp long EGFP exon 1, encoding only 53 AAs, is co-expressed in cells expressing the ORFeus-IS reporter. This transcript presents less than one-quarter of the whole EGFP protein sequence (SFig. 3) no longer displaying most of the immunogenic epitopes recognized by commercially available EGFP antibodies. We carried out a minimal necessary modification of the traditional mouse ORFeus reporter by shifting the position of the intron in the EGFP coding sequence and did not introduce or remove any additional nucleotide sequences. This modification enables the selective IHC-based detection of the full-length fluorescent EGFP protein, which appears exclusively in cells and their progeny that have undergone ORFeus retrotransposition, separately from the polypeptide encoded by EGFP exon 1, present in all ORFeus reporter-expressing cells (SFig. 3). This is essential for the reliable single-cell level detection of reporter retrotransposition by IHC.

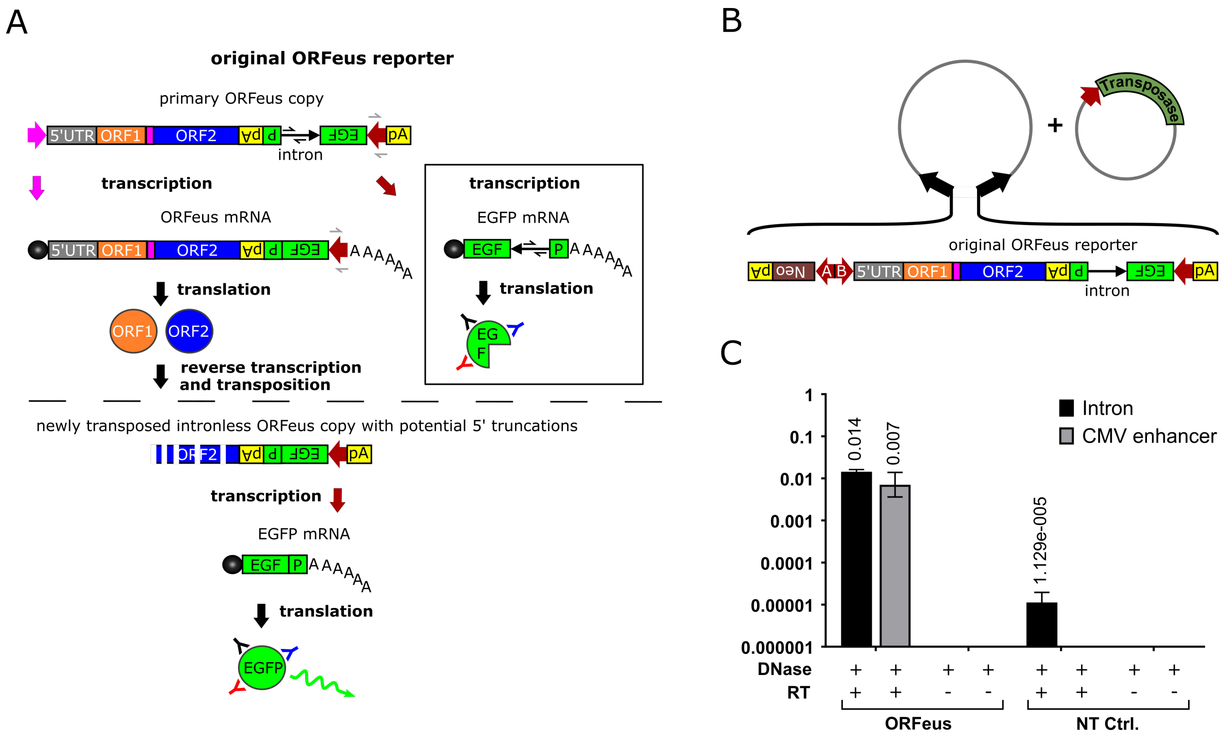


**Fig. S1. The operating principle of ORFeus reporter with the resulting transcripts and their RTqPCR-based quantitation**

**(A)** Schematic representation of the working principle of the ORFeus reporter. The original ORFeus transgene expresses a sense-oriented L1 transcript and an antisense-oriented reporter transcript. The antisense reporter transcript boxed in black is potentially capable of expressing the polypeptide encoded by the first exon of EGFP. Following successful retrotransposition of the L1 transcript, the newly generated genomic ORFeus copies produce an antisense reporter transcript that is intron-free and capable of expressing the full-length fluorescent EGFP protein. Pink arrow, sense promoter; red arrow, antisense promoter; pink rectangle, inter-ORF spacer; black ball, mRNA cap structure; black chevron arrows, intron-specific RT-qPCR primers; gray chevron arrows, CMV enhancer-specific RT-qPCR primers; black, red and blue Ys, antibodies. (**B**) Schematic representation of the *Sleeping Beauty* (SB) transposon-based cloning platform for the stabile expression of ORFeus type reporters in mammalian cell culture. Black arrows, SB transposon inverted terminal repeats; red arrows, promoters; pink rectangle, inter-ORF spacer. (**C**) Monitoring the amount of the L1 and the reporter transcripts in HT1080 cells after stable SB transposon-based delivery of the ORFeus element. RNA samples were collected from cultured non-transfected (NT) cells and from ORFeus transfected cells after G418 selection, then tested using an intron- and a CMV enhancer-specific RT-qPCR assay. Results were normalized to measurements of the ribosomal protein L27 (Rpl27) transcript as input control and data were presented on a logarithmic scale as the geometric mean ± standard deviation (SD) (n = 3) (see Supplementary Materials 2 for individual data values and statistics).

A

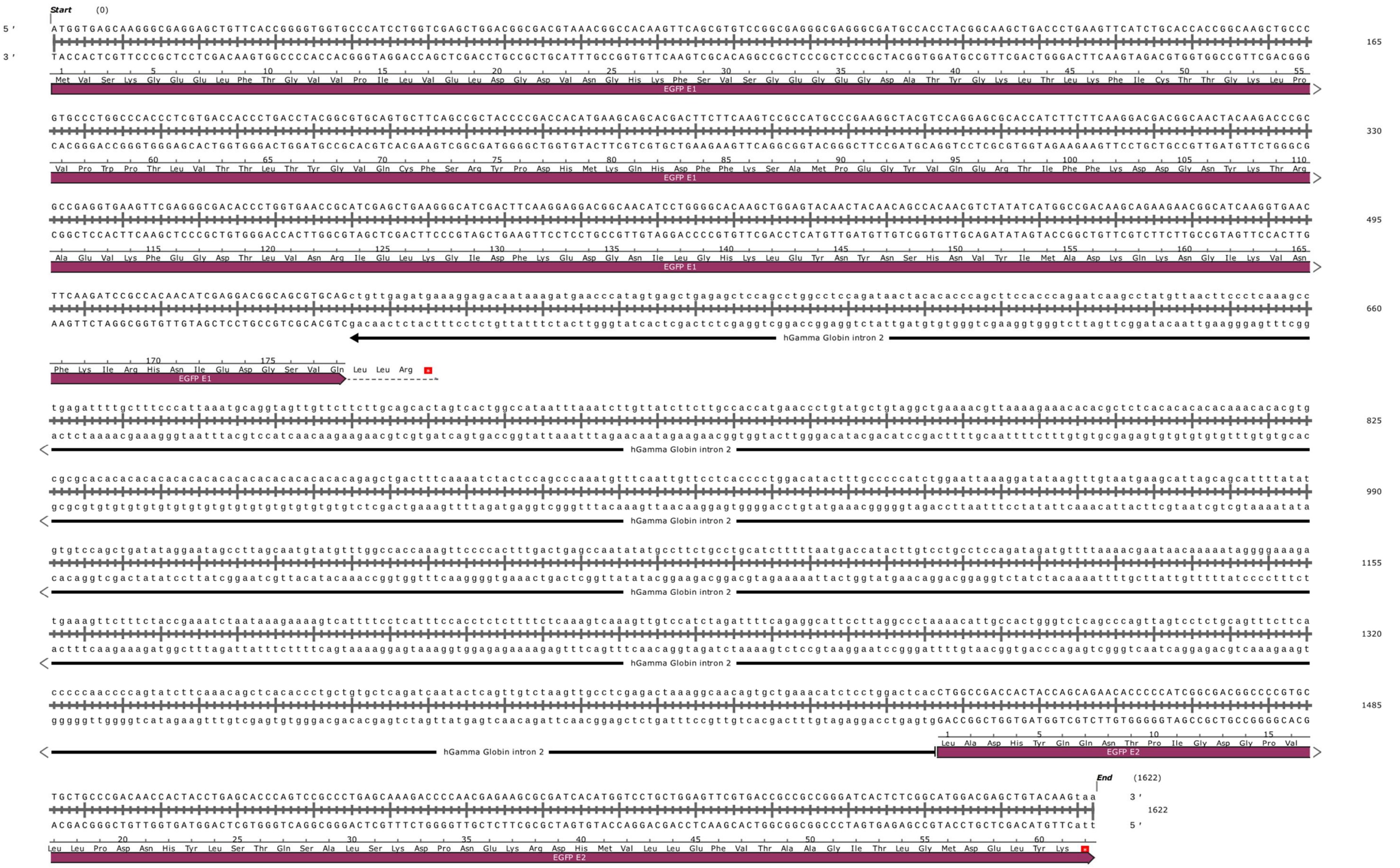
Start (0)
EGFP E1
hGamma Globin intron 2
EGFP E2
End (1622)
165
330
495
660
825
990
1155
1320
1485
1622

B

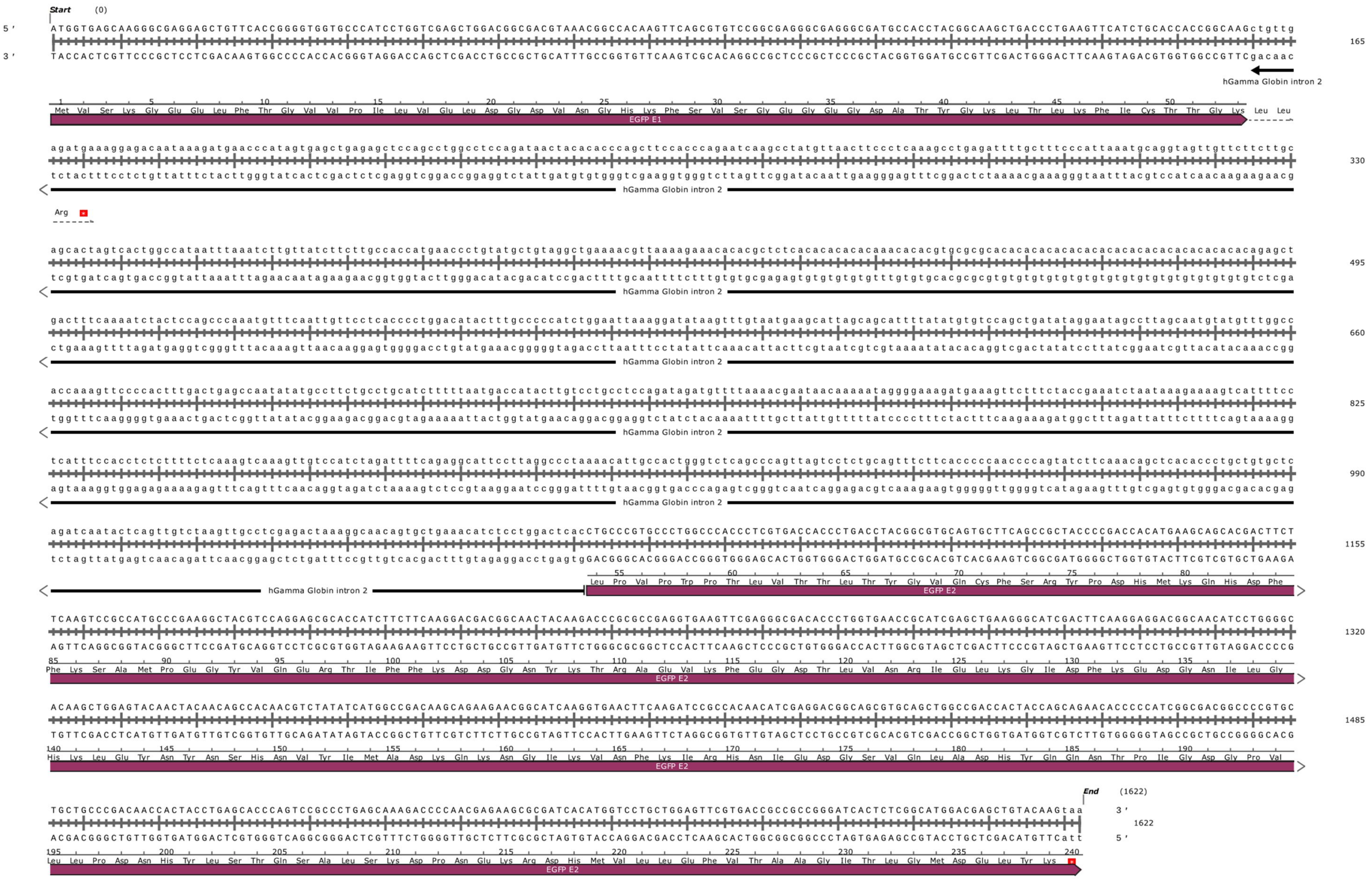

Start (0)
5'
3'
165
330
495
660
825
990
1155
1320
1485
1622
End (1622)
hGamma Globin intron 2
EGFP E1
EGFP E2

**Fig. S2. Nucleotide sequence of a section of the EGFP retrotransposition indicator cassette present in the ORFeus and ORFeus-IS vectors**

**(A)** Nucleotide sequence of the EGFP exons and the modified human γ-globin intron found in the original ORFeus vectors. **(B)** Nucleotide sequence of the EGFP exons and the modified human γ-globin intron in the ORFeus-IS vectors. Solely the position of the intron was changed within the EGFP coding sequence.

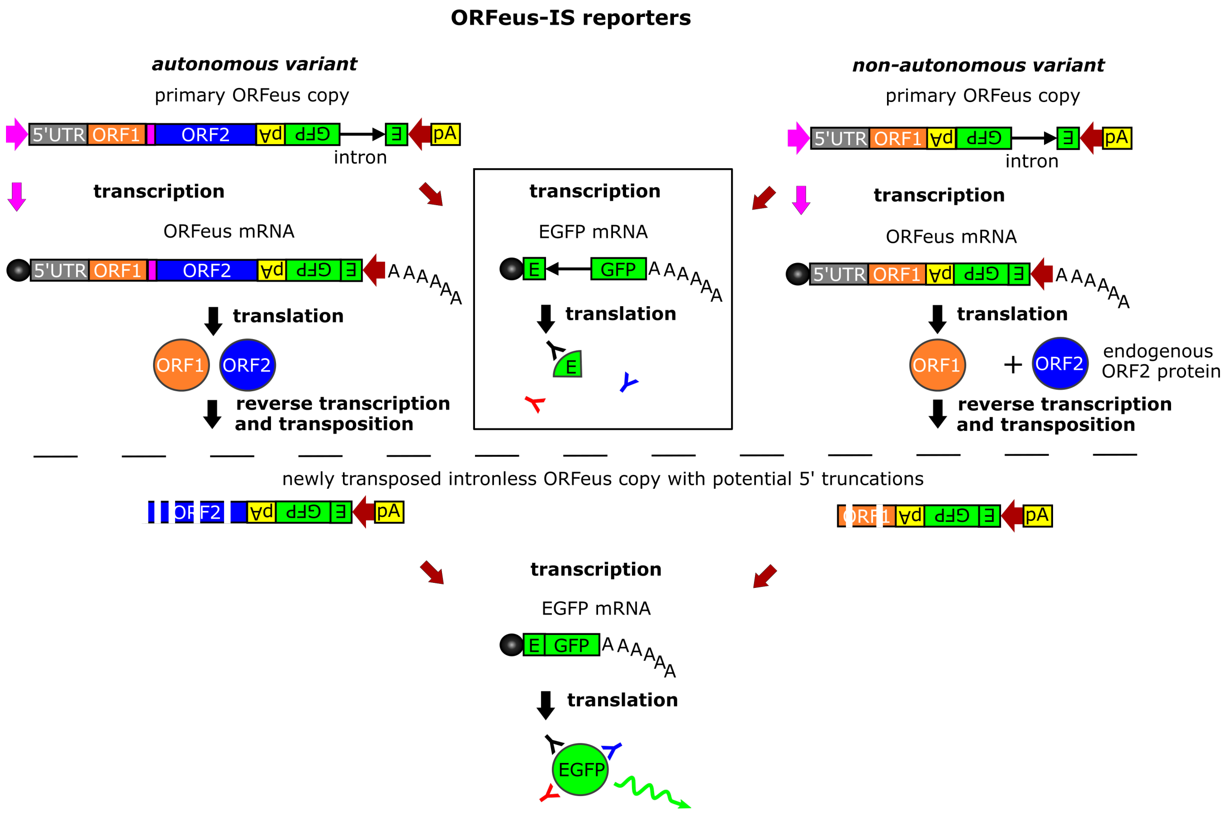


**Fig. S3. Schematic representation of the structure and operating principle of the ORFeus-IS and NA ORFeus-IS elements.**

The ORFeus-IS transgenes express a sense-oriented L1 transcript and an antisense-oriented reporter transcript. However, the antisense reporter transcript (boxed in black) is only capable of expressing a shortened 159 bp long EGFP exon 1. Following successful retrotransposition of the L1 transcript, the newly generated genomic ORFeus-IS copies produce an antisense reporter transcript that is intron-free and capable of expressing the full-length fluorescent EGFP protein. In the case of the NA ORFeus-IS reporter variant, the L1-ORF2p protein is provided by endogenous L1 copies present in the host cell genome. Pink arrow, sense promoter; red arrow, antisense promoter; pink rectangle, inter-ORF spacer; black ball, mRNA cap structure; black, red and blue Ys, antibodies.

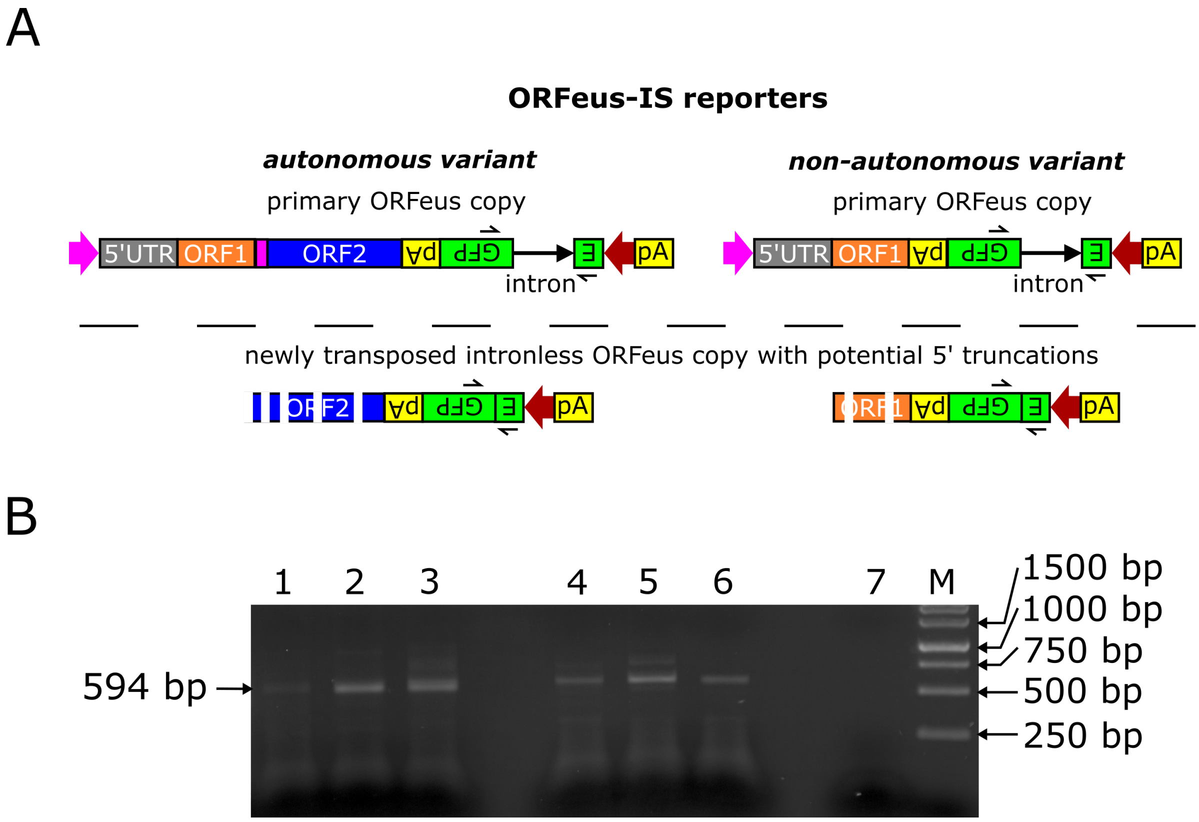


**Fig. S4. Confirmation of ORFeus-IS reporter retrotransposition at the DNA level in the liver of mice**

**(A)** Schematic representation of the structure of the original ORFeus-IS and NA ORFeus-IS transgenes and their descendants that have undergone splicing and retrotransposition (shown only at the DNA level). The positions of the primers used for the intron-spanning PCR assay are also indicated. The size of the PCR product generated by the primer pair is 1496 bp in the original transgenes and 594 bp in the secondary DNA copies that have undergone retrotransposition. Pink arrow, sense promoter; red arrow, antisense promoter; pink rectangle, inter-ORF spacer; black chevron arrows, intron-spanning PCR primers. (**B**) Result of the intron-spanning PCR assay that is able to detect loss of the EGFP intron in newly transposed L1 reporter copies. Liver DNA samples were prepared 3 months after hydrodynamic injection and tested by the intron-spanning PCR assay. PCR products were analyzed by agarose gel electrophoresis. The PCR assay confirmed the presence of intron-free retrotransposed ORFeus copies in livers carrying both ORFeus-IS and NA ORFeus-IS elements. Lanes 1-3, DNA samples prepared from ORFeus-IS bearing livers; lanes 4-6, DNA samples prepared from NA ORFeus-IS bearing livers; lane 7, no template control; M, DNA size marker.

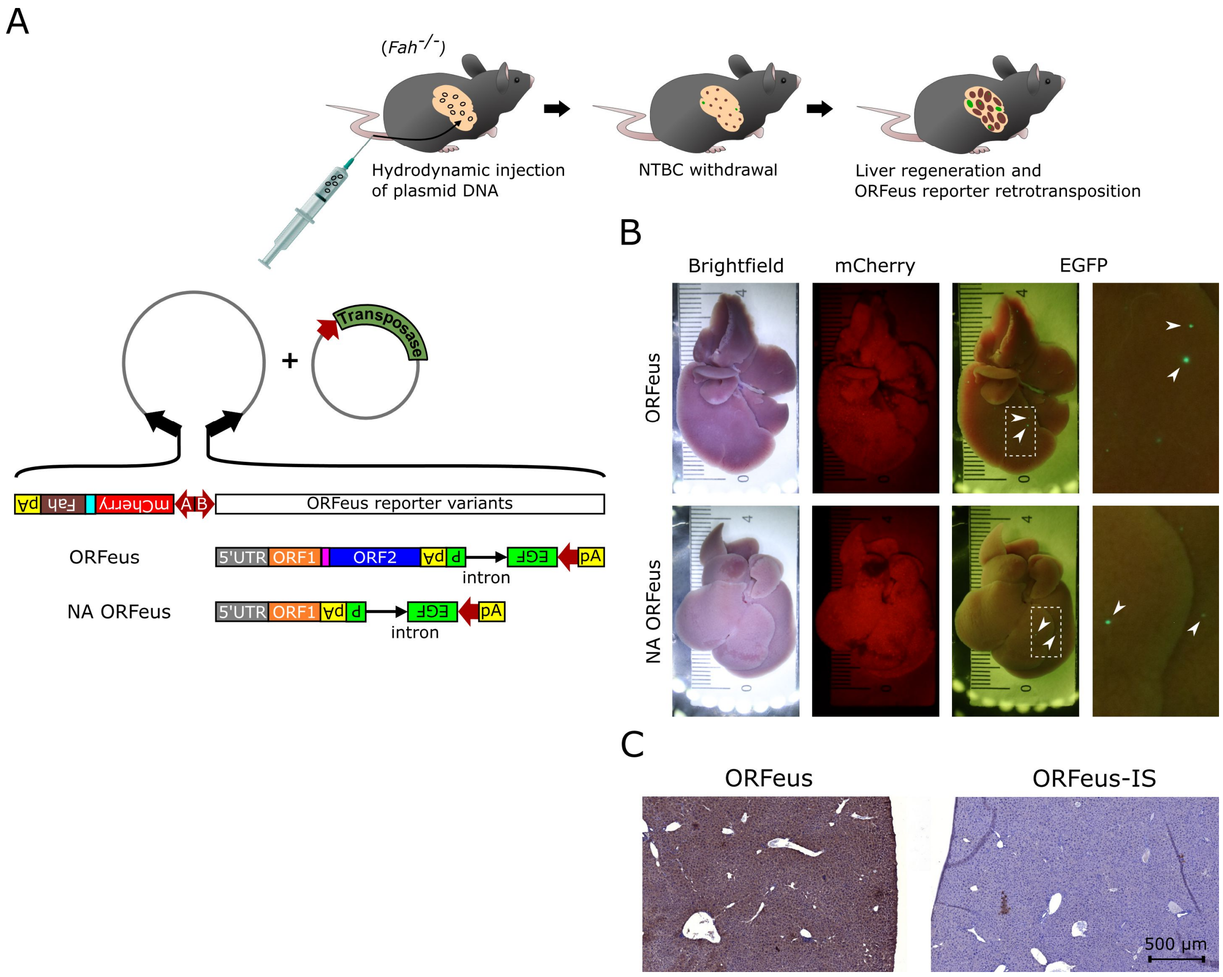


**Fig. S5.** ***In vivo*** **characterization of the traditional ORFeus element and its non-autonomous variant in the mouse liver**

**(A)** Schematic representation of the structure of the *PiggyBac* (PB) transposon-based cloning platform and animal treatments. Black arrows, PB transposon inverted terminal repeats; red arrows, promoters; pink rectangle, inter-ORF spacer. **(B)** Brightfield and fluorescence stereomicroscopic images of the liver of *Fah*$^{-/-}$ mice 3 months after the intrahepatic delivery of ORFeus and NA ORFeus elements. (**C**) EGFP immunostainings of liver sections from *Fah*$^{-/-}$ mice, 3 months after hydrodynamic injection of ORFeus and ORFeus-IS elements. The immunostainings were performed in parallel, side by side.
