## Supplementary material for "Intrahepatic reporter assay reveals leaky somatic blockade of L1 retrotransposition in mice": Individual data values and statistics

**Fig. 1B**

| | ORFeus | | | ORFeus-IS | | | NA ORFeus | | | NA ORFeus-IS | | |
|---|---|---|---|---|---|---|---|---|---|---|---|---|
| Repetitions | 1 | 2 | 3 | 1 | 2 | 3 | 1 | 2 | 3 | 1 | 2 | 3 |
| Values | 79,314 | 78,964 | 66,62 | 21,124 | 20,56 | 20,816 | 0,024 | 0,022 | 0,036 | 0,022 | 0,002 | 0,012 |
| Mean | 74,97 | | | 20,83 | | | 0,02733 | | | 0,012 | | |

ORFeus-IS vs ORFeus
Unpaired t test with Welch's correction
P value 0,0058 **

NA ORFeus vs ORFeus
Unpaired t test with Welch's correction
P value 0,0031 **

NA ORFeus-IS vs ORFeus-IS
Unpaired t test with Welch's correction
P value <0,0001 ****

NA ORFeus-IS vs NA ORFeus
Unpaired t test with Welch's correction
P value 0,1067 **ns**

### Fig. 2E

| | ORFeus-IS | | | NA ORFeus-IS | | |
|---|---|---|---|---|---|---|
| Animal | 1 | 2 | 3 | 1 | 2 | 3 |
| Values | 0,064005 | 0,087333 | 0,045373 | 0,018844 | 0,019985 | 0,027255 |
| Mean | 0,06557 | | | 0,02203 | | |

NA ORFeus-IS vs ORFeus-IS
Unpaired t test with Welch's correction
P value 0,0637 **ns**

### Fig. 2F

| | ORFeus-IS | | | NA ORFeus-IS | | |
|---|---|---|---|---|---|---|
| Animal | 1 | 2 | 3 | 1 | 2 | 3 |
| Values | 80,43748 | 88,8215 | 78,76995 | 56,13022 | 46,74301 | 44,61192 |
| Mean | 82,68 | | | 49,16 | | |

NA ORFeus-IS vs ORFeus-IS
Unpaired t test with Welch's correction
P value 0,0022 **

### Fig. 2G

| | | Number of EGFP positive colonies in colony classes defined according to cell count, per 1 million hepatocytes | | | | | | | |
|---|---|---|---|---|---|---|---|---|---|
| | | ORFeus-IS | | | | NA ORFeus-IS | | | |
| Animal | | 1 | 2 | 3 | **Avg** | 1 | 2 | 3 | **Avg** |
| colony classes (cell counts) | **1-10** | 64,09862 | 71,26376 | 64,70389 | **66,68876** | 54,12557 | 44,83514 | 40,15073 | **46,37048** |
| | 1 | 31,42089 | 19,62336 | 14,06606 | | 20,04651 | 20,03272 | 14,87064 | |
| | 2 | 8,797849 | 19,62336 | 15,47267 | | 12,0279 | 8,585451 | 11,89651 | |
| | 3 | 6,284178 | 8,262465 | 14,06606 | | 10,02325 | 3,815756 | 5,948256 | |
| | 4 | 8,797849 | 6,196849 | 5,626425 | | | 4,769695 | | |
| | 5 | 3,770507 | 2,065616 | 1,406606 | | 2,004651 | 2,861817 | 1,487064 | |
| | 6 | 1,256836 | 8,262465 | 8,439638 | | 2,004651 | 2,861817 | | |
| | 7 | | 3,098425 | 2,813213 | | 2,004651 | 0,953939 | 1,487064 | |
| | 8 | 2,513671 | | 1,406606 | | 6,013952 | 0,953939 | 2,974128 | |
| | 9 | | 4,131233 | 1,406606 | | | | 1,487064 | |
| | 10 | 1,256836 | | | | | | | |
| | **11-20** | 8,797849 | 7,229657 | 12,65946 | **9,562321** | 2,004651 | 3,815756 | 2,974128 | **2,931512** |
| | **21-30** | 2,513671 | 3,098425 | | **1,870699** | | 0,953939 | | **0,31798** |
| | **31-40** | 2,513671 | 1,032808 | | **1,18216** | | | | |
| | **41-50** | | 1,032808 | 1,406606 | **0,813138** | | | | |
| | **51-60** | | 1,032808 | | **0,344269** | | | 1,487064 | **0,495688** |
| | **61-70** | 1,256836 | 2,065616 | | **1,107484** | | | | |
| | **71-80** | 1,256836 | 1,032808 | | **0,763215** | | | | |
| | **81-90** | | 1,032808 | | **0,344269** | | | | |

**SFig. 1C**

| | ORFeus | | | ORFeus, no RT | | | NT Ctrl. | | | NT Ctrl., no RT | | |
|---|---|---|---|---|---|---|---|---|---|---|---|---|
| Repetitions | 1 | 2 | 3 | 1 | 2 | 3 | 1 | 2 | 3 | 1 | 2 | 3 |
| Intron specific assay | | | | | | | | | | | | |
| Values (normalized to Rpl27) | 0,015734 | 0,012604 | 0,014989 | | | | | | | 0,000015 | 0,000006 | 0,000016 |
| Mean | 0,014442333 | | | | | | | | | 1,23333E-05 | | |
| Geo. mean | 0,014378273 | | | | | | | | | 1,12924E-05 | | |
| CMV enhancer specific assay | | | | | | | | | | | | |
| Values (normalized to Rpl27) | 0,003331 | 0,012007 | 0,008851 | | | | | | | | | |
| Mean | 0,008063 | | | | | | | | | | | |
| Geo. mean | 0,007074034 | | | | | | | | | | | |
